# Simulating multi-substrate diffusive transport in 3-D tissues with BioFVM

**DOI:** 10.1101/035709

**Authors:** Samuel H. Friedman, Ahmadreza Ghaffarizadeh, Paul Macklin

**Author notes:** Research supported by University of Southern California (USC) Center for Applied Molecular Medicine (CAMM), the Breast Cancer Research Foundation, the NIH (5U54CA143907, 1R01CA180149), and the USC James H. Zumberge Research and Innovation Fund. S. H. F., A. G., and P. M. are all with the USC CAMM, 2250 Alcazar St, Rm 240, Los Angeles, CA, 90089. P. M. is the corresponding author. S. H. F. is. A. G. is.

## Abstract

To simulate the spatiotemporal distribution of chemical compounds, we present BioFVM, an open-source reaction-diffusion equation solver using finite volume methods with motivation for biological applications. With various numerical solvers, we can simulate the interaction of dozens of compounds, including growth substrates, drugs, and signaling compounds in 3-D tissues, with cells by treating them as various source/sink terms. BioFVM has linear computational cost scalings and demonstrates first-order accuracy in time and second-order accuracy in space. Beyond simulating the transport of drugs and growth substrates in tissues, the ability to simulate dozens of compounds should make 3-D simulations of multicellular secretomics feasible.

## I. Introduction to the type of problem in cancer

Tissues are filled with various chemical compounds, including signaling and other factors that regulate how cells move, grow, and die, depending on the concentration and/or gradient of any and all of these compounds. In order for cancer cells to survive and grow, they need to obtain oxygen and other nutrients released from blood vessels, and change phenotype based upon signaling factors released by other cells and the vasculature. These chemical substances move through tissues by diffusion, and are impacted by uptake by tumor and other cells and reaction terms (e.g., decay). (In some tissues, advection is also dominant; these effects are not considered in this version.) These same transport processes can be used to model how chemotherapeutic drugs reach their intended targets: susceptible cancer cells. By performing computational simulations of the movement of dozens of various chemical substances and of cellular chemical uptake and secretion rates, we can test hypotheses that can control the overall growth of cancer cells and enable 3-D simulations of multicellular secretomics.

Simulating this motion requires solving a system of reaction-diffusion equations for a vector 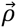 of substrates:

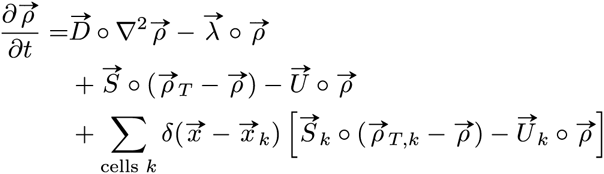

where **D** and **λ** are the diffusion and decay coefficients, **S** and **U** are the bulk source and uptake terms, and **S***_k_* and **U***_k_* are cell-centered source and uptake terms. The **o** symbol denotes the Hadamard (element-wise) product between two vector terms, *δ* is the Dirac delta function, and ***ρ***_T_ represents target densities of the substrates.

We present here BioFVM [1], an open-source software package written in C++ to efficiently solve this system of equations on multicore desktop computer and single supercomputer nodes. Using the finite volume method (FVM) [2], this software fills a niche and focuses on the biological transport processes of interest to cancer modelers. BioFVM is intended to work in concert with multicellular modeling packages (e.g., Chaste [3], CompuCell3D [4], PhysiCell [5]) to simulate how cells change their phenotypes as they interact with other cells and the biochemical microenvironment. Combining BioFVM with an agent-based code enables a user to simulate the growth, survival, and death of not only cancer cells, but of multicellular processes in general.

BioFVM is open source under the Apache 2.0 license. It can be downloaded for free at:

http://BioFVM.MathCancer.orgORhttp://BioFVM.sf.net.

A series of tutorials is included with every download. The method is described in detail in [1].

## II. Illustrative Results of Application of Methods

BioFVM obtains first-order accuracy in time, second-order accuracy in space, and linear scaling of cost in terms of each of the number of substrates, voxels, and time steps [1]. Simulating additional substrates does not significantly increase the computational cost of a simulation. Fig. 1 demonstrates the computational cost scaling when increasing the number of substrates from 1 to 100.

**Figure 1:**
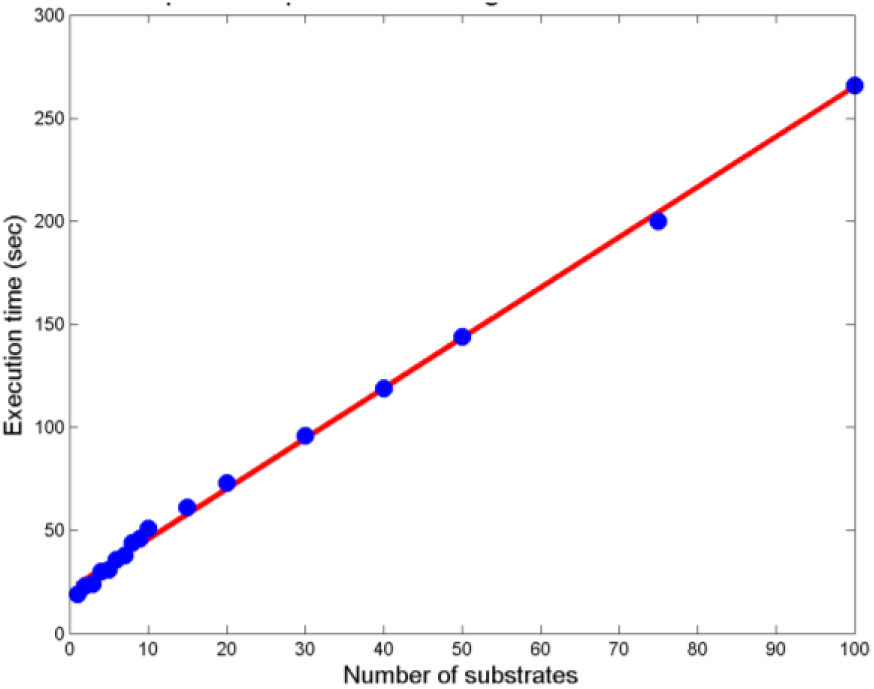
Number of substrates vs. execution time (seconds) for a simulation with otherwise identical conditions.

Fig. 2 shows an example simulating oxygen and glucose transport in a liver lobule. We marked central veins in histological images of liver tissues, estimated the corresponding lobular geometry, generated sinusoids, added hepatocytes, and imported tissue into BioFVM with a 5 mm^3^ computational domain (5 mm × 5 mm × 200 μm). We simulated oxygen and glucose release by the portal triads, diffusion, and uptake by hepatocytes. The desktop simulation used 625,000 voxels (20 μm resolution) with 900,000 cells. See [1] for further examples and test results.

**Figure 2:**
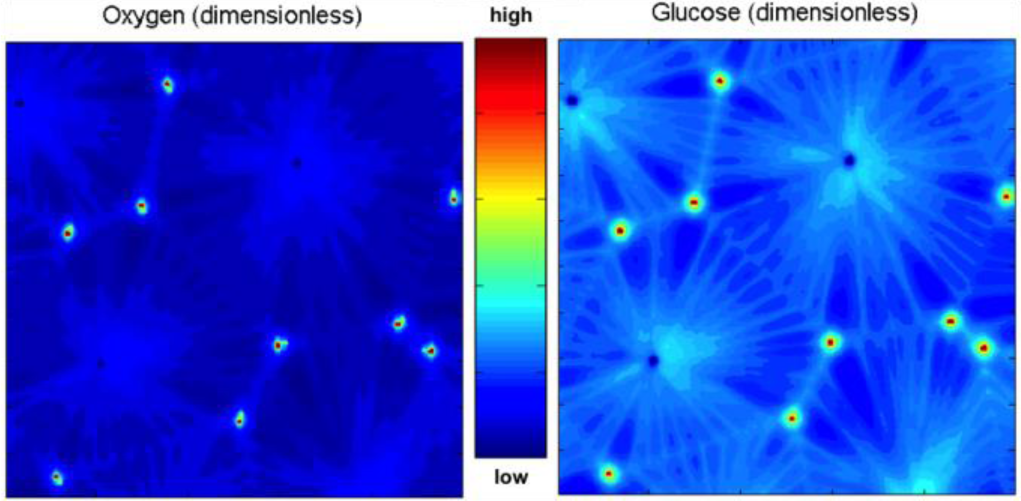
Simulating transport of oxygen (left) and glucose (right) in virtual liver lobules (hexagonal shapes). Nondimensionalized substrate values increase from blue to red.

## III. Quick Guide to the Methods

We discretize the spatial simulation domain into voxels to solve the reaction-diffusion equation with the finite volume method. BioFVM currently supports uniform Cartesian meshes, where the FVM reduces to a finite difference scheme; support for non-Cartesian Voronoi meshes is expected in a future release.

We several techniques to speed up numerical computations. We use operator splitting [6, 7] to solve the bulk source/sink terms first, the cellular source/sink terms second, and diffusion/decay terms third. The splitting is *O*(Δt) accurate and stable when the individual solvers are first-order accurate and stable [6]. Moreover, it simplifies development, allows efficient solvers to be tailored to each term, and holds the door open to new solvers (e.g., advection) without significant change to the overall code.

We solve the bulk and cell-centered source/sink terms using the (1^st^-order, stable) backwards Euler scheme. In each time step, the source/sink terms are independent across space. Thus, parallelize their solutions using OpenMP. See [1] for the full details of the discretization.

We divide the 3-D diffusion/decay terms into a series of 1-D equations, one for each dimension [6]. After this splitting, a strip of voxels (e.g. all voxels with the same *y_j_*· and *z_k_* values, but different *x* values) is independent of any other strip; discretizing the diffusion and decay terms yields a tridiagonal linear system, which can be directly by using the Thomas algorithm [8]. Because each *x* strip is independent, we can distribute and run many instances of the Thomas solver across the processor cores with OpenMP, allowing us to easily parallelize the *x* diffusion problem. We solve along each of the *y* and *z* dimensions similarly.

To further optimize the code, we used some computational techniques that better utilize modern computer architecture, trading memory for computational speed wherever possible. Rather than solving the PDE for each chemical substrate sequentially (which requires 10× more computational time to solve for 10 substrates than solving for one substrate), we discretized the solvers to operate on entire vector of substrates simultaneously, thus taking advantage of SIMD and similar hardware optimizations on modern CPUs. We used BLAS operations (e.g. axpy) on C++ vectors to accelerate computations to avoid hidden memory allocations/copies [9]. Due to these optimizations, we found that simulating 10 substrates requires approximately 2.6× more computational time than a single substrate [1]. Because the diffusion and decay coefficients are assumed constant, we could pre-compute the forward sweep and part of the back substitution steps for the Thomas solvers, allowing further improvement in the computational speed. [1].

## Acknowledgment

We thank the USC Center for Applied Molecular Medicine for generous resources. This work was supported by the Breast Cancer Research Foundation for support, the National Institutes of Health (Physical Sciences Oncology Center grant 5U54CA143907 for Multi-scale Complex Systems Transdisciplinary Analysis of Response to Therapy (MC-START), and 1R01CA180149), and the USC James H. Zumberge Research and Innovation Fund.

## References

[1] A. Ghaffarizadeh, S. H. Friedman, and P. Macklin. (2015, December 12, 2015). BioFVM: an efficient, parallelized diffusive transport solver for 3-D biological simulations. Bioinformatics. Available: http://dx.doi.org/10.1093/bioinformatics/btv730

[2] R. Eymard, T. Gallouët, and R. Herbin, “The finite volume method,” in Handbook for Numerical Analysis. vol. 7, P. G. Ciarlet and J. L. Lions, Eds., ed: North Holland, 2000, pp. 713–1018.

[3] G. R. Mirams, C. J. Arthurs, M. O. Bernabeu, R. Bordas, J. Cooper, A. Corrias, Y. Davit, S. J. Dunn, A. G. Fletcher, D. G. Harvey, M. E. Marsh, J. M. Osborne, P. Pathmanathan, J. Pitt-Francis, J. Southern, N. Zemzemi, and D. J. Gavaghan, “Chaste: an open source C++ library for computational physiology and biology,” PLoS Comput Biol, vol. 9, p. e1002970, 2013.

[4] M. H. Swat, G. L. Thomas, J. M. Belmonte, A. Shirinifard, D. Hmeljak, and J. A. Glazier, “Multi-scale modeling of tissues using CompuCell3D,” Methods Cell Biol, vol. 110, pp. 325–66, 2012.

[5] (2015-). PhysiCell Project Website (Note: To be submitted and open sourced in 2016). Available: http://PhysiCell.MathCancer.org

[6] G. I. Marchuk, “Splitting and alternating direction methods,” in Handbook of Numerical Analysis. vol. Volume 1, ed: Elsevier, 1990, pp. 197–462.

[7] B. Seibold, “18.336 Numerical Methods for Partial Differential Equations Spring 2009,” Massachusetts Institute of Technology: MIT OpenCouseWare2009.

[8] L. H. Thomas, “Elliptic problems in linear difference equations over a network,” Columbia University, New York, New York1949.

[9] L. S. Blackford, J. Demmel, J. Dongarra, I. Duff, S. Hammarling, G. Henry, M. Heroux, L. Kaufman, A. Lumsdaine, A. Petitet, R. Pozo, K. Remington, and R. C. Whaley, “An updated set of basic linear algebra subprograms (BLAS),” ACM Trans. Math. Softw., vol. 28, pp. 135–151, 2002.

